# Distinct virus-specific regulation of RNA synthesis across genome segments by thogotovirus polymerases: insights from Oz virus and Dhori virus

**DOI:** 10.64898/2026.03.31.715722

**Authors:** Tofazzal Md Rakib, Rena Mashimo, Lipi Akter, Hiroshi Shimoda, Yudai Kuroda, Hiromichi Matsugo, Yusuke Matsumoto

## Abstract

Thogotoviruses are a group of tick-borne, six-segmented, negative-sense single-stranded RNA viruses. These viruses encode an RNA-dependent RNA polymerase that recognizes promoter sequences located at the genomic termini to initiate RNA synthesis. The 5′ and 3′ ends of the genome bind to the polymerase and function as a promoter. Outside the catalytic center, they base-pair with each other to form a double-stranded RNA structure. This structure is referred to as the distal duplex and plays an important role in RNA synthesis. In this study, we investigated how the RNA sequence of the distal duplex influences polymerase activity using minigenome systems of two thogotoviruses, Oz virus (OZV) and Dhori virus (DHOV). Each virus exhibits distinct activities among its six segments. In OZV, one determinant of these differences is the base pair at positions 5′12 and 3′11 within the distal duplex, where promoter activity varies depending on whether the base pair is G:C or A:U. In contrast, the DHOV polymerase is not affected by this difference. These results indicate that, even within the genus Thogotovirus, viruses differ in whether they possess a mechanism that modulates promoter activity based on subtle sequence differences within the distal duplex. Furthermore, phylogenetic analysis and comparison of promoter sequences suggest that thogotoviruses can be divided into groups that do or do not regulate intersegment promoter activity via the base pair at positions 5′12 and 3′11.

**Highlights:**

- Minigenome systems of Oz virus and Dhori virus reveal segment-specific differences in promoter activity
- The distal duplex sequence modulates RNA synthesis in a virus-dependent manner
- The base pair at positions 5′12/3′11 determines promoter activity in Oz virus but not in Dhori virus
- Thogotoviruses can be divided into groups that do or do not regulate promoter activity via distal duplex sequence variation at positions 5′12/3′11

## Introduction

Members of the genus *Thogotovirus* (family Orthomyxoviridae) are tick-borne RNA viruses (Hubálek et al., 2014). This genus includes Thogoto virus (THOV), Dhori virus (DHOV), Oz virus (OZV), Bourbon virus, and several related viruses. Thogotoviruses infect mammals, causing febrile illness and neurological symptoms. THOV and DHOV are known to infect humans as well as domestic and wild animals, and human infections can result in febrile disease or encephalitis (Butenko et al., 1987; Filipe et al., 1985; Hubálek and Rudolf, 2012). OZV was first isolated from ticks in Japan in 2018 (Ejiri et al., 2018), and one human infection has been reported to date (Osawa et al., 2025). In that case, the patient developed fever and systemic symptoms and ultimately died of viral myocarditis. Thus, thogotoviruses represent a group of zoonotic viruses of public health importance.

These viruses possess a segmented, negative-sense single-stranded RNA genome consisting of six segments (segment 1 - 6), which encode PB2, PB1, PA, glycoprotein (GP), nucleoprotein (NP), and matrix protein (M), respectively. The viral RNA genome is encapsidated by NP and is recognized by the RNA-dependent RNA polymerase (RdRp) complex composed of PA, PB1, and PB2, which mediates genome replication and mRNA transcription (Dick et al., 2024; Xue et al., 2024).

Structural analyses of the THOV polymerase have shown that the 5′ and 3′ termini of the viral genome bind to the RdRp complex and form a partially double-stranded RNA structure through complementary base pairing in regions slightly internal to the termini (Akter et al., 2025; Xue et al., 2024). This structure, termed the distal duplex, is thought to play an important role in the initiation of viral RNA synthesis.

We previously established a minigenome system for OZV and reported that RNA synthesis activity differs among genome segments (Akter et al., 2025). However, it remains unclear whether this segment-specific regulation of RNA synthesis is unique to OZV or represents a mechanism shared among thogotoviruses. In the present study, we established a minigenome system for DHOV, a virus closely related to OZV, and compared RNA synthesis activities among segments.

## Materials and Methods

### Cells

BHK/T7-9 cells (Ito et al., 2003) were cultured in Dulbecco Modified Eagle’s Medium with 10% fetal calf serum and penicillin/streptomycin. Cells were cultured at 37°C in 5% CO_2_.

### Plasmid construction

The Nano luciferase (Nluc)-expressing OZV (Akter et al., 2025) and DHOV minigenome plasmids were constructed using the pCAGGS plasmid backbone. The Nluc gene was flanked by the 3’ and 5’ untranslated regions (UTR) of the OZV EH8 strain genome (GenBank accession numbers: segment 1; NC_040730.1, segment 2; NC_040731.1, segment 3; NC_040732.1, segment 4; NC_040735.1, segment 5; LC320127.2, and segment 6; LC320128.2) and DHOV Dhori/1313/61 genome (GenBank accession numbers: segment 1; NC_034261.1, segment 2; NC_034254.1, segment 3; NC_034255.1, segment 4; NC_034262.1, segment 5; NC_034256.1, and segment 6; NC_034263.1). The minigenome is set under the control of the RNA polymerase-II promoter, and the transcript expressed as a negative sense RNA is cleaved at both ends by a hammerhead ribozyme (Rz) and a hepatitis delta virus (HDV) Rz (Beaty et al., 2017). Both OZV and DHOV PA, PB1, PB2 and NP genes, which are encoded in segments 3, 2, 1 and 5 respectively, are set into the multiple cloning sites of pCAGGS vector. All plasmids were generated through custom gene synthesis services by GenScript (GenScript Japan, Tokyo, Japan). A pCAGGS-derivative plasmid for the expression of firefly luciferase (Fluc) was also constructed (Ashida et al., 2024).

### OZV and DHOV Nluc minigenome assay

The OZV and DHOV Nluc minigenome assay was performed in BHK/T7-9 cells cultured in 12-well plates (1 × 10^5^ cells/well). Plasmids; OZV/DHOV-Nluc (0.4 μg), pCAGGS-NP (0.4 μg), -PB1 (0.4 μg), -PB2 (0.4 μg) and -PA (0.4 μg), and Fluc (0.05 μg) were transfected using XtremeGENE HP (Merck, Darmstadt, Germany) according to the manufacturer’s instructions. At 48 hour post-transfection, the cells were lysed with Passive Lysis Buffer and the Nluc and Fluc activities were measured using the Nano-Glo Dual-Luciferase Reporter Assay System (Promega, Madison, WI, USA) according to the manufacturer’s instructions. The mRNAs of Fluc and Nluc are transcribed by the cellular polymerase and viral polymerases, respectively. Since the Fluc values are assumed to remain constant across all samples, the Nluc values were normalized to the Fluc values. These normalized Nluc values were then used to calculate relative values for each experiment.

### Statistical analysis

A two-tailed unpaired Student’s t-test was employed to compare the means of the two independent groups. The test was performed in R using the t.test() function. The t-statistic and *p*-value were calculated, and a *p*-value of less than 0.05 indicated a statistically significant difference between the groups.

## Results

### Promoter activity of OZV and DHOV minigenomes

We previously reported, using an OZV minigenome system, that RNA synthesis activity differs among genome segments. Specifically, segments 1, 2, and 3 exhibited high activity, whereas segments 4, 5, and 6 showed relatively lower activity (Akter et al., 2025). In OZV, differences in base pairs within the distal duplex were correlated with RNA synthesis activity among segments. Specifically, when the base pair at positions 5′12 and 3′11 was G:C, high activity was observed (segments 1, 2, and 3), whereas when it was A:U, lower activity was observed (segments 5 and 6) (Fig. 1A). In segment 4, although this position contained a G:C base pair, the activity was relatively low.

**Figure 1.**
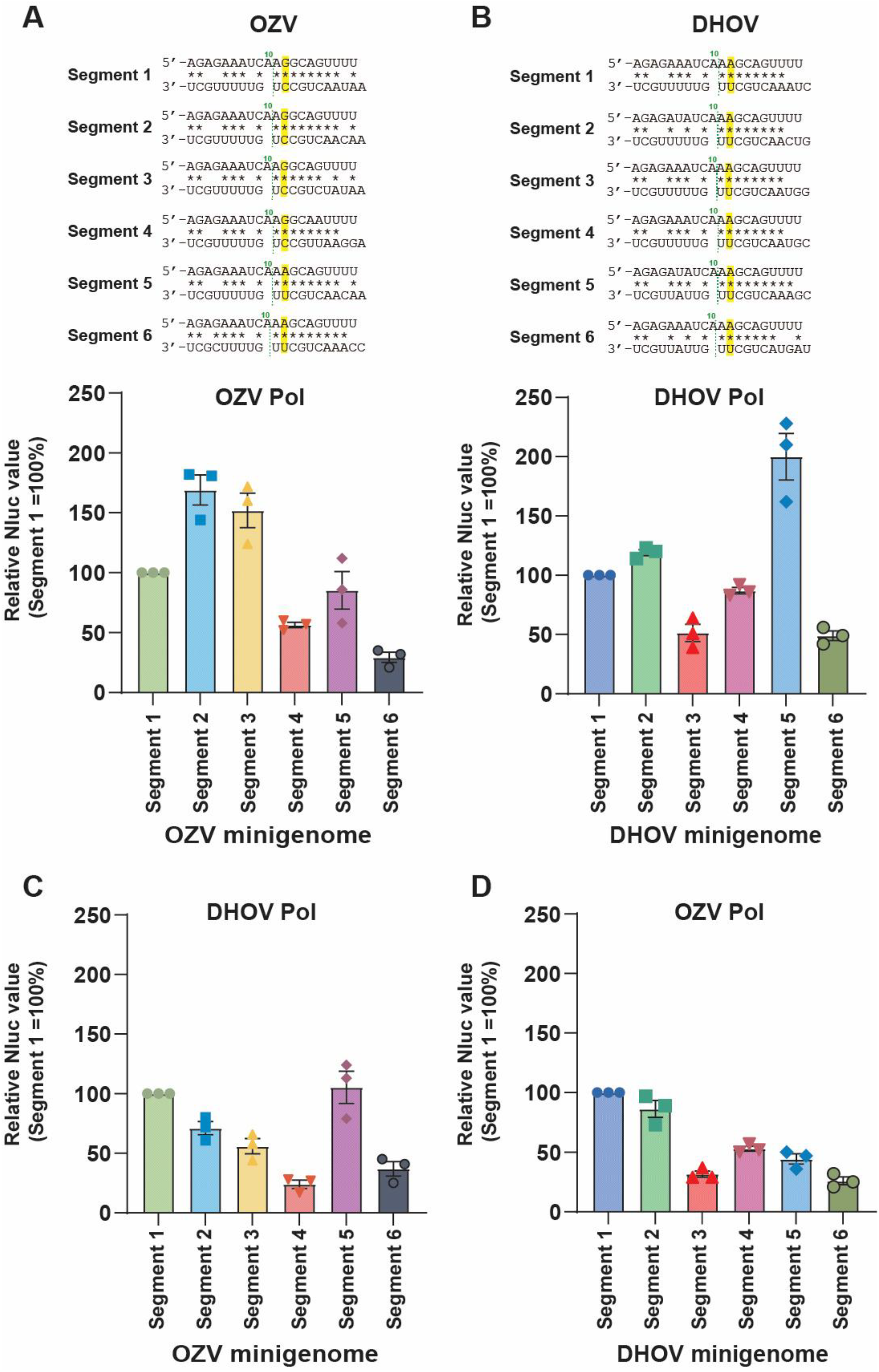
Recognition of viral promoters by OZV and DHOV polymerases. **(A, B)** Promoter structures of OZV **(A)** and DHOV **(B)** segments 1–6. Complementarity between nucleotides at positions 1–9 of the 5′ and 3′ genomic termini, as well as between nucleotides from position 11 onward at the 5′ end and from position 10 onward at the 3′ end, was analyzed. The base pair between the 12th nucleotide at the 5′ end and the 11th nucleotide at the 3′ end is highlighted in yellow. **(C, D)** Minigenome activities measured using OZV or DHOV polymerase components (PA, PB1, PB2, and NP) with the indicated viral segments. **(C)** DHOV polymerase with OZV segments 1–6. **(D)** OZV polymerase with DHOV segments 1–6. For panels **(A–D)**, relative Nluc values are shown (segment 1 = 100%). Data represent the mean ± standard deviation from three independent experiments. Pol indicates polymerase.

To examine whether such segment-specific promoter activity is also present in other thogotoviruses, we established a minigenome system for DHOV. Minigenomes containing the promoter sequences of DHOV segments 1–6 were constructed, and their activities were measured using the DHOV polymerase components (NP, PA, PB1 and PB2) (Fig. 1B). Segment 5 (encoding NP) exhibited the highest activity, followed by segment 2 (PB1), segment 1 (PB2), and segment 4 (GP). However, the differences among segments, except for segment 5, were smaller than those observed in OZV (Fig. 1A). Analysis of the DHOV promoter sequences revealed that the base pair at positions 5′12 and 3′11 was A:U in all segments. Moreover, the distal duplex region, consisting of at least seven base pairs (5′11–17 nt / 3′10–16 nt), was identical among all six segments (Fig. 1B).

### Differential recognition of OZV and DHOV promoters by polymerases

Next, we examined how the polymerase components (PA, PB1, PB2, and NP) of OZV and DHOV recognize promoter sequences from each virus using a minigenome system. When the DHOV polymerase was used, activities with OZV promoters differed from those obtained with the OZV polymerase on OZV promoters (Fig. 1A, C). When the OZV polymerase was used, activities with DHOV promoters differed from those obtained with the DHOV polymerase on DHOV promoters (Fig. 1B and D). Importantly, the relatively higher activities observed for OZV promoters 2 and 3 compared with OZV promoter 1 were not detected when the DHOV polymerase was used.

We then introduced substitutions in the distal duplex of the OZV promoters of segment 5 and 6, changing the base pair at positions 5′12 and 3′11 from A:U to G:C. As previously reported (Akter et al., 2025), when the OZV polymerase was used, this substitution resulted in an approximately fourfold increase in activity for segment 5 (Fig. 2A: OZV pol) and an approximately threefold increase for segment 6 (Fig. 2B: OZV pol). In contrast, when the DHOV polymerase was used, no significant change in polymerase activity was observed for segment 5, whereas a statistically significant but modest decrease in activity was observed for segment 6 (Fig. 2A and B: DHOV pol).

**Figure 2.**
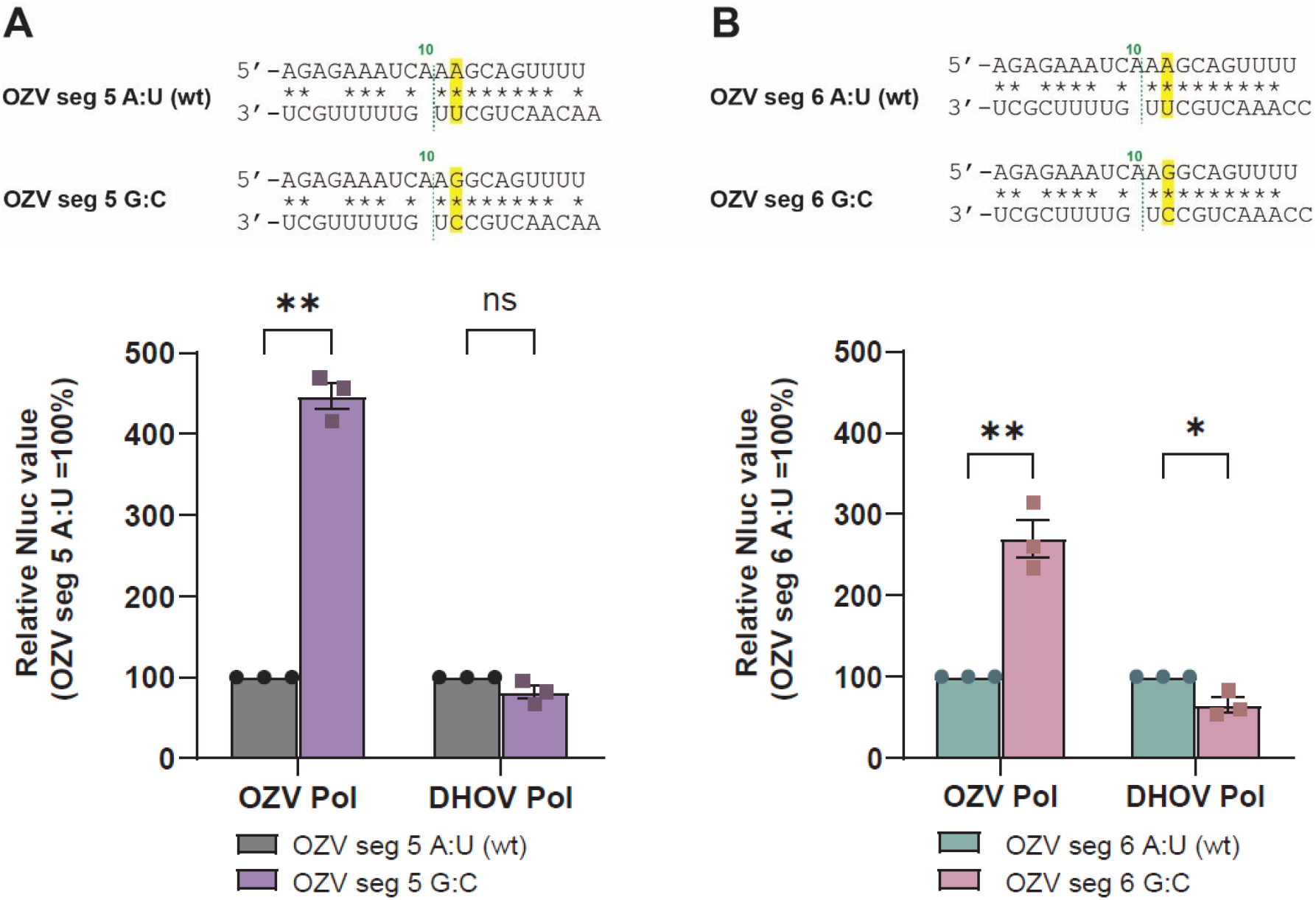
Effect of the 5′12/3′11 base pair in the OZV distal duplex on promoter activity. **(A, B)** Minigenome activities of OZV segments 5 **(A)** and 6 **(B)** using A:U wild-type (wt) or G:C mutant constructs, measured with OZV or DHOV polymerase components (PA, PB1, PB2, and NP). Relative Nluc values are shown (A:U (wt) = 100%). Data represent the mean ± standard deviation from three independent experiments. ns, not significant; \**p* < 0.05; \*\**p* < 0.01 (two-tailed unpaired Student’s t-test).

We further introduced the same base-pair substitution into DHOV minigenomes of segments 5 and 6, changing the base pair at positions 5′12 and 3′11 from A:U to G:C. When the OZV polymerase was used, this substitution increased activity by approximately eightfold for segment 5 and approximately twentyfold for segment 6 (Fig. 3A and B: OZV pol). However, when the DHOV polymerase was used, no change in activity was observed (Fig. 3A and B: DHOV pol).

**Figure 3.**
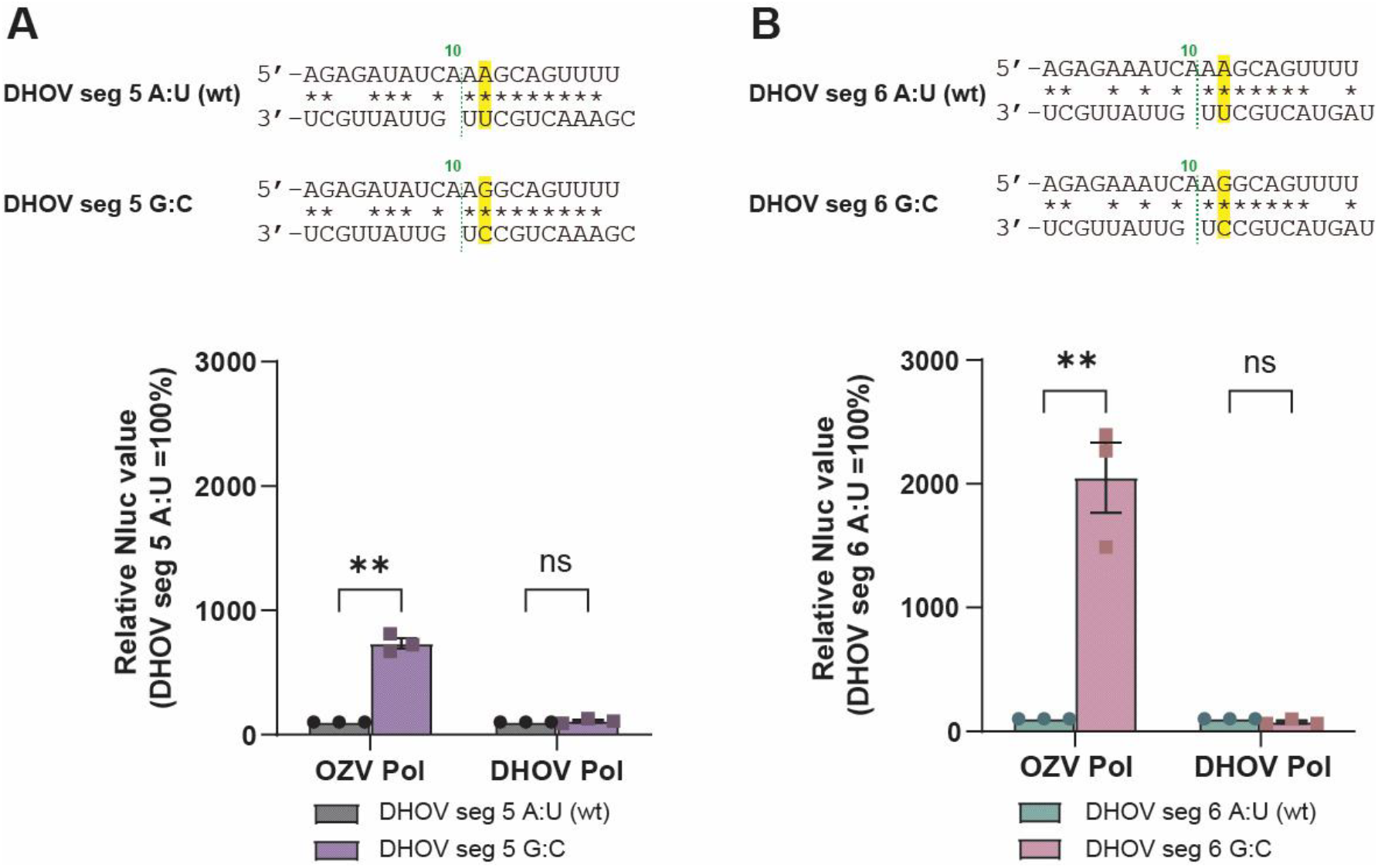
Effect of the 5′12/3′11 base pair in the DHOV distal duplex on promoter activity. **(A, B)** Minigenome activities of DHOV segments 5 **(A)** and 6 **(B)** using A:U wt or G:C mutant constructs, measured with OZV or DHOV polymerase components (PA, PB1, PB2, and NP). Relative Nluc values are shown (A:U wt = 100%). Data represent the mean ± standard deviation from three independent experiments.

### Effect of distal duplex mutations in OZV Segment 4

Although OZV segment 4 contains a G:C base pair at positions 5′12 and 3′11 in the distal duplex, it exhibits relatively low activity with the OZV polymerase (Fig. 1A) (Akter et al., 2025). In this segment, the base pair at positions 5′16 and 3′15 is A:U, which differs from that in other segments (Fig. 1A). To examine the effect of this base pair, we constructed a minigenome in which this position was changed to G:C and measured its activity. When the OZV polymerase was used, this substitution resulted in an approximately threefold increase in activity (Fig. 4: OZV pol). When the DHOV polymerase was used, a similar approximately threefold increase in activity was observed (Fig. 4: DHOV pol).

**Figure 4.**
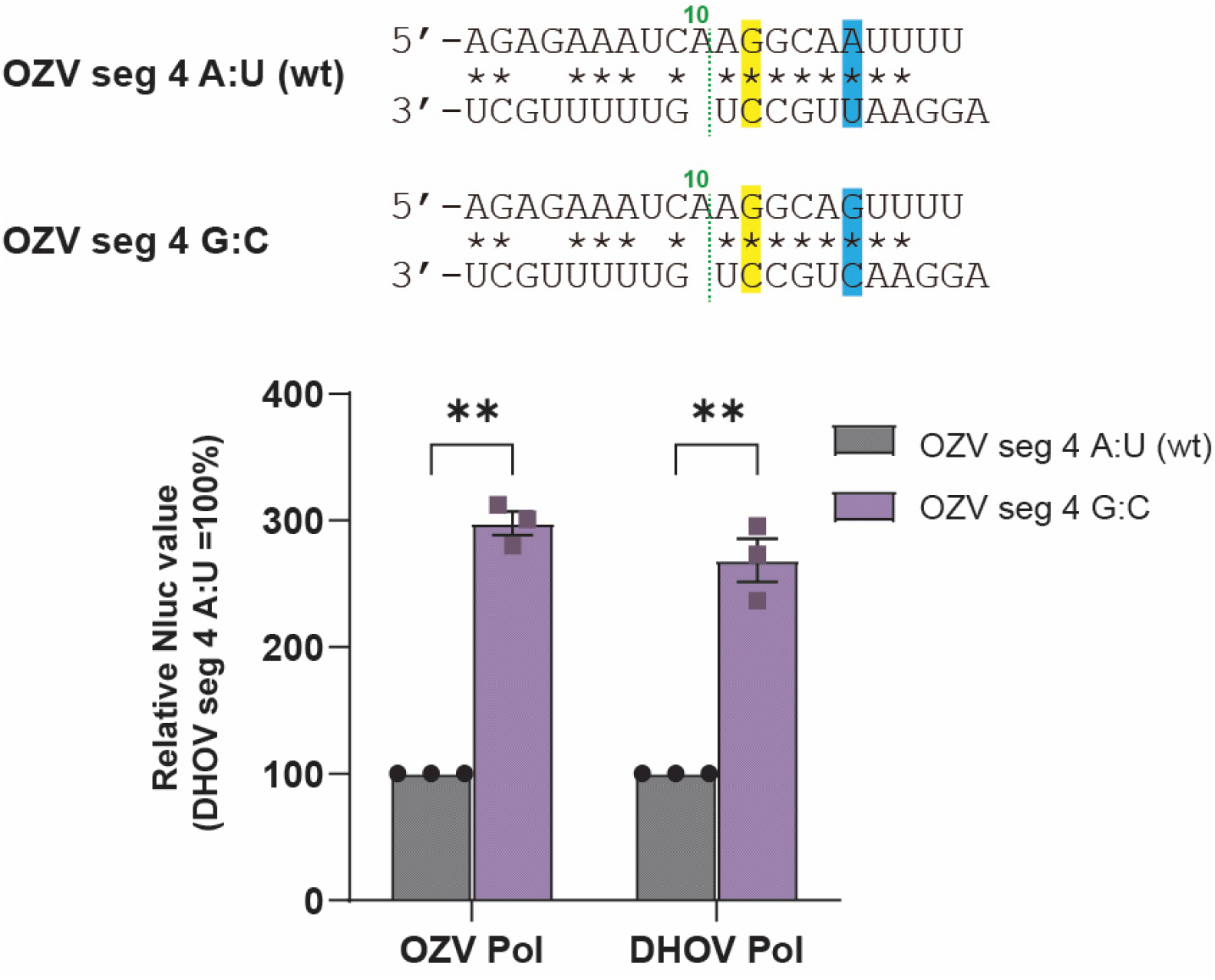
Effect of the 5′16/3′15 base pair in the distal duplex of OZV segment 4 on promoter activity. Minigenome activities of OZV segments 4 using A:U wt or G:C mutant constructs, measured with OZV or DHOV polymerase components (PA, PB1, PB2, and NP). Relative Nluc values are shown (A:U wt = 100%). Data represent the mean ± standard deviation from three independent experiments. The base pair between the 16th nucleotide at the 5′ end and the 15th nucleotide at the 3′ end is highlighted in blue.

### Comparison of promoter sequences among thogotoviruses

Promoter sequences were compared among thogotoviruses whose complete terminal sequences for all six segments are available in the GenBank database at the National Center for Biotechnology Information (NCBI). The analysis included Upolu virus, Aransas Bay virus, Thogoto virus, Oz virus, Dhori virus, and Bourbon virus (Fig. 5) (Briese et al., 2014). Among these viruses, two patterns were observed with respect to the base pair at positions 5′12 and 3′11 in the distal duplex. In Dhori virus and Bourbon virus, all six segments contained an A:U base pair at this position. In contrast, in Upolu virus, Aransas Bay virus, Thogoto virus and Oz virus, both A:U and G:C base pairs were present among segments.

**Figure 5.**
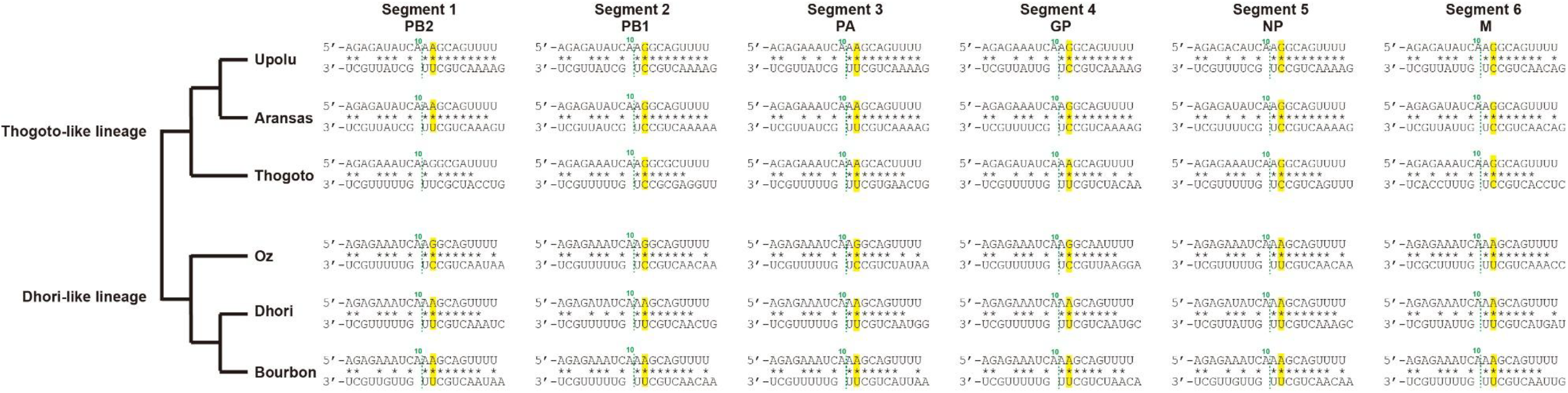
Comparison of distal duplex base pairs among thogotoviruses. Promoter structures of segments 1–6 from Upolu virus, Aransas Bay virus, Thogoto virus, Oz virus, Dhori virus, and Bourbon virus. Complementarity between nucleotides at positions 1–9 of the 5′ and 3′ genomic termini, as well as between nucleotides from position 11 onward at the 5′ end and from position 10 onward at the 3′ end, was analyzed. The base pair between the 12th nucleotide at the 5′ end and the 11th nucleotide at the 3′ end is highlighted in yellow. A phylogenetic tree based on the PB1 gene indicates that these viruses are divided into two lineages, the Thogoto-like and Dhori-like lineages. In DHOV and Bourbon virus, the 5′12/3′11 base pair is A:U in all segments, whereas in Upolu virus, Aransas Bay virus, Thogoto virus and Oz virus, both A:U and G:C base pairs are observed among segments.

A phylogenetic tree based on the PB1 gene indicates that these six viruses can be divided into two major lineages: the Thogoto-like lineage (Upolu virus, Aransas Bay virus, and Thogoto virus) and the Dhori-like lineage (Oz virus, Dhori virus, and Bourbon virus) (Fig. 5) (Fuchs et al., 2022; Kosoy et al., 2015). Within the Dhori-like lineage, Oz virus is unique in that the base pair at positions 5′12 and 3′11 varies among segments (A:U or G:C), whereas Dhori virus and Bourbon virus consistently possess an A:U base pair at this position in all segments. In contrast, viruses belonging to the Thogoto-like lineage exhibit variation at this position, with both A:U and G:C base pairs observed among segments.

### Exchangeability of polymerase components between OZV and DHOV

The thogotovirus polymerase is composed of a complex of PA, PB1, PB2, and NP. To examine the exchangeability of these components between OZV and DHOV, NP, PA, PB1, and PB2 derived from each virus were expressed in various combinations, and their activities were assessed using a minigenome assay. Using the DHOV segment 1 minigenome, a total of 16 combinations were tested (Fig. 6). Clear polymerase activity was observed only when PA, PB1, and PB2 were derived from the same virus. In contrast, NP was interchangeable between the two viruses. However, when DHOV PA, PB1, and PB2 were combined with OZV NP, polymerase activity was markedly reduced.

**Figure 6.**
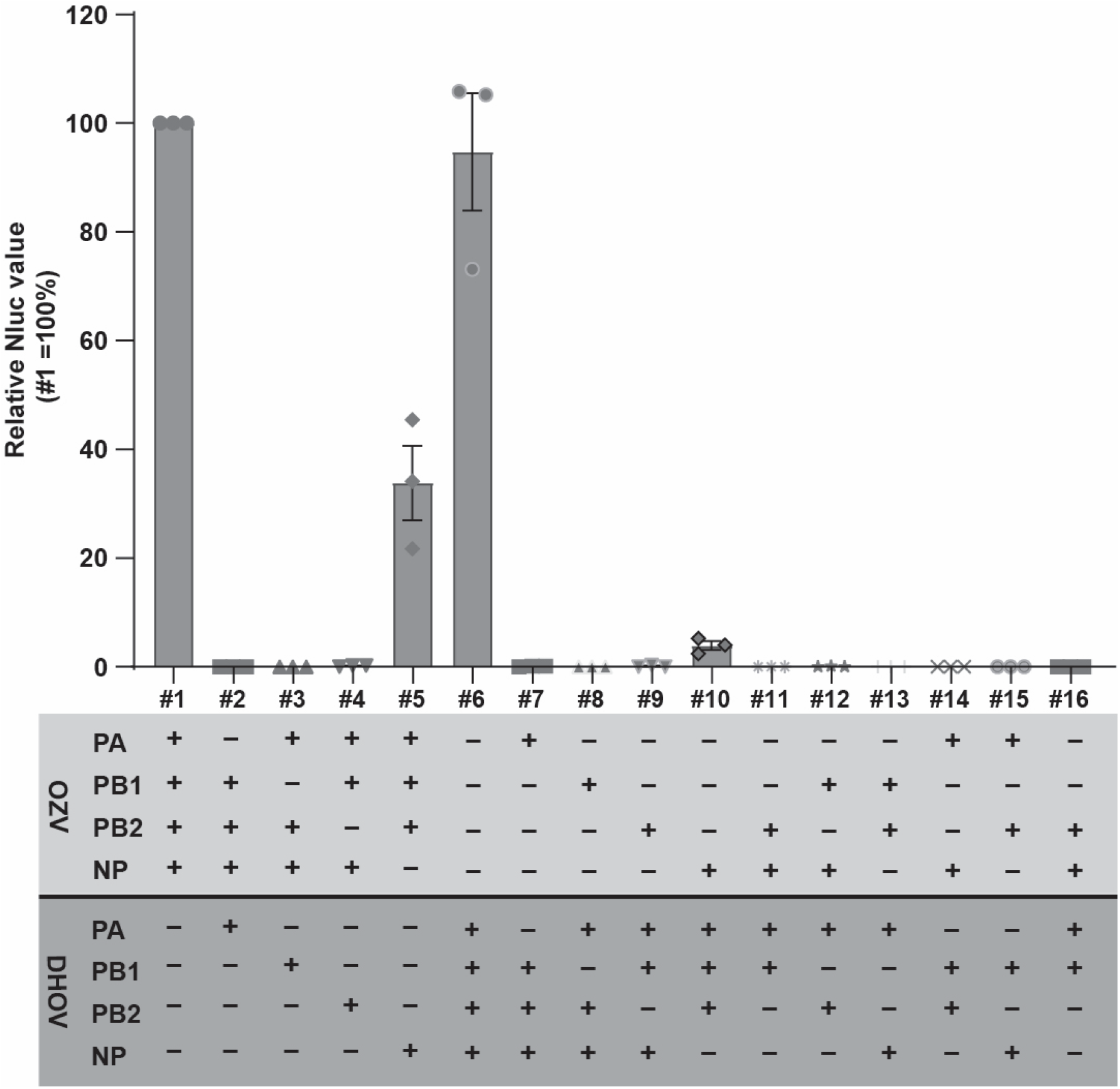
Exchangeability of polymerase components between OZV and DHOV. Minigenome activities of the DHOV segment 1 promoter were measured using OZV or DHOV polymerase components (PA, PB1, PB2, and NP) in all 16 possible combinations of subunit exchange. Relative Nluc values are shown (#1 = 100%). Data represent the mean ± standard deviation from three independent experiments.

## Discussion

Promoter activity in OZV is influenced by the base pair at positions 5′12 and 3′11 within the distal duplex, with G:C supporting higher activity than A:U (Fig. 2A and B: OZV pol) (Akter et al., 2025). In contrast, changing this base pair had no effect on DHOV polymerase activity (Fig. 2A and B: DHOV pol), indicating that DHOV polymerase is largely insensitive to variation at this position. These findings indicate that polymerase–promoter recognition varies among thogotoviruses, with virus-specific differences in position-specific sensitivity within the distal duplex.

This observation can be interpreted in the context of the promoter structure revealed in the THOV polymerase complex (Xue et al., 2024). Structural analyses of the THOV polymerase have revealed a characteristic arrangement of the viral RNA promoter. During RNA synthesis initiation, nucleotides 1–8 at the 3′ end of the vRNA are positioned in the polymerase active site. Nucleotides 1–10 at the 5′ end form a hook structure through intra-strand base pairing in the 5′ hook binding pocket within the polymerase, as seen in other segmented negative-strand RNA viruses (Malet et al., 2023; Pflug et al., 2017). Outside of the active site, nucleotides from the 5′ and 3′ termini form a complementary RNA duplex, referred to as the distal duplex, which is also seen in polymerases conformation of diverse segmented negative-strand RNA viruses (Neriya et al., 2022; Pflug et al., 2014). This duplex functions as a structural scaffold that positions the 3′ end of the vRNA for proper entry into the active site, thereby facilitating the initiation of RNA synthesis (Xue et al., 2024). In the THOV structure, the distal duplex consists of at least seven complementary base pairs. Comparison of the terminal sequences of OZV indicates that this region forms at least six base pairs, corresponding to nucleotides 5′11–16 and 3′10–15 (Akter et al., 2025). The present OZV results suggest that the identity of the 5′12/3′11 base pair (G:C or A:U) may influence transcription initiation by altering the stability of this RNA duplex. In contrast, the lack of a similar effect in DHOV suggests that its polymerase is functionally less sensitive to differences in distal duplex stability.

Comparison of promoter sequences among thogotoviruses available in public databases revealed two distinct patterns (Fig. 5). In some viruses, the base pair at positions 5′12 and 3′11 varies (G:C or A:U) among genome segments, whereas in others it is identical (A:U) across all segments, as observed in DHOV and Bourbon virus. Taken together, thogotovirus polymerases can be broadly classified into two types: those whose promoter activity is sensitive to the GC/AU state of the 5′12/3′11 base pair within the distal duplex and those that are largely insensitive to this difference. The phylogenetic tree (Fig. 5) indicates that members of the genus *Thogotovirus* are broadly divided into two groups, the Thogoto-like and Dhori-like lineages (Fuchs et al., 2022). OZV, DHOV, and Bourbon virus belong to the Dhori-like group. Within this lineage, DHOV and Bourbon virus are more closely related to each other, whereas OZV occupies a slightly more distant position (Fig. 5). If the GC/AU-insensitive polymerase represents the predominant type among Dhori-like viruses, OZV may represent an exceptional case within this lineage from the perspective of polymerase–promoter evolution.

In addition, we confirmed that in both OZV and DHOV, as observed for segment 4, the base pair at positions 5′16 and 3′15 also influences promoter activity (Fig. 4). Importantly, in DHOV, the base pair at positions 5′12/3′11 had no effect on activity, whereas changes at positions 5′16/3′15 altered polymerase activity. These observations suggest that the positions within the distal duplex that influence promoter activity differ between viruses, at least between OZV and DHOV.

Thogotoviruses possess segmented RNA genomes and therefore have the potential to undergo reassortment when two viruses infect the same cell (Davies et al., 1987; Jones et al., 1987). In this study, we showed that the polymerases of OZV and DHOV are capable of recognizing promoter sequences derived from the other virus (Fig. 1), suggesting that a certain level of compatibility exists between heterologous viral segments at the level of promoter recognition. However, polymerase component exchange experiments demonstrated that clear activity was observed only when PA, PB1, and PB2 were derived from the same virus (Fig. 6). Thus, even if reassortment occurs at the genome segment level, the formation of a functional polymerase complex may require specific combinations of polymerase subunits. This observation contrasts with influenza A and B viruses, in which certain polymerase components from different viruses can sometimes function together (Iwatsuki-Horimoto et al., 2008). Thus, even among segmented viruses within the family *Orthomyxoviridae*, the exchangeability of polymerase complexes and the mechanisms of promoter recognition appear to vary among viruses.

## Conclusion

This study demonstrates that, in thogotoviruses, even a single base pair within the distal duplex that is likely located outside direct contact with the polymerase can modulate promoter activity. Moreover, sensitivity to this base pair differs among viruses. Identifying the polymerase determinants responsible for this differential sensitivity between OZV and DHOV will provide further mechanistic insight into position-specific promoter recognition by thogotovirus polymerases.

## Acknowledgements

We thank Dr. Naoto Ito (Gifu University) for providing the BHK/T7-9 cells. We also thank Dr. Kyoko Tsukiyama-Kohara (Kagoshima University) and members of her laboratory for their support with the biological experiments.

